# Directing conservation action for the Critically Endangered Philippine Eagle to mitigate mining impacts and maximize indigenous land management

**DOI:** 10.1101/2025.05.19.654818

**Authors:** Luke J. Sutton, Jayson C. Ibañez, Dennis I. Salvador, Andrei Von Mariano C. Tirona, Guiller S. Opiso, Tristan Luap P. Senarillos, Kristian J. Suetos, Rowell L. Taraya, Christopher J.W. McClure

## Abstract

As habitat destruction intensifies due to expanding human infrastructure, balancing biodiversity conservation with resource extraction has become a global challenge. Thus, quantifying the extent and location of proposed mining operations is key to mitigating the impacts on threatened species. This issue is particularly acute in the biodiversity hotspots of southeast Asia, where rapid economic growth needs to be balanced with sustainable conservation management of the remaining tropical forest. The Philippine Eagle (*Pithecophaga jefferyi*), a critically endangered tropical forest raptor endemic to the Philippines, faces increasing threats from mining activities that destroy and fragment its tropical forest habitat. Here, we integrate Species Distribution Modelling with gap and hotspot analysis to assess the spatial overlap between Philippine Eagle nest habitat and mining concessions across a protected area network on Mindanao, where the largest population remains. Using a landscape-scale SDM built with remote sensing covariates and eagle occurrence data, we identified high-suitability nest habitats and projected these into the Eastern Mindanao Biodiversity Corridor (EMBC). Hotspot analysis revealed that 41 % of the total mining concessions area contained high-suitability nest habitat, highlighting significant conservation risks. Additionally, 46 % of indigenous ancestral domains contain high-suitability nest habitat, emphasizing the importance of indigenous land management in safeguarding eagle habitats. Our gap analysis demonstrated that 35 % of the EMBC protects high-suitability nest habitat, which here represents the remaining montane tropical forest. To mitigate mining impacts, we propose targeted nest surveys in high-risk areas, implementation of mining moratoriums near critical nesting zones, and strengthening indigenous and protected area land management. By integrating SDMs with spatial analysis, we provide a framework for directing conservation efforts to balance resource extraction with the preservation of the Philippine Eagle’s habitat. Our findings offer crucial insights for policy development and land-use planning to protect this iconic species and its threatened habitat.

## Introduction

Mining is one of the most significant anthropogenic activities driving habitat loss (Sonter *et al*. 2018). The extraction of minerals and other earth-based resources involves land clearance, soil excavation, and water diversion, which leads to habitat destruction and pollution of natural systems. These changes often create unsuitable conditions for species, driving some to local extinction and reducing biodiversity overall (Sonter *et al*. 2018; Betts *et al*. 2021). Addressing the impacts of mining on biodiversity requires integrated conservation strategies, including land-use planning and enforcement of environmental policies, to balance resource extraction with biodiversity protection.

As a key global biodiversity hotspot, the Philippines is home to numerous endemic and threatened species, many of which depend on the remaining tropical forest for survival (Myers *et al*. 2000). Mining extractions in the Philippines are often carried out in ecologically sensitive areas, which accelerate deforestation, fragment habitats, and alter the physical and chemical properties of the environment, posing severe threats to wildlife populations (BirdLife International 2017). The extraction of minerals involves clearing forests, excavating land, and generating waste that can pollute rivers and streams (WWF 2015). These activities not only degrade ecosystems but also displace species, reducing available habitat and isolating populations (Laurance *et al*. 2014). Mining operations also create noise, vibration, and human presence that can disturb sensitive species, particularly those with specific habitat or breeding requirements (Paguntalan & Jakosalem 2008).

The Philippine Eagle (*Pithecophaga jefferyi*) is an evolutionary distinct tropical forest raptor (McClure *et al*. 2023) and one of the most threatened birds globally, currently classified as ‘Critically Endangered’ on the IUCN Red List (BirdLife International 2018). The Philippine Eagle is endemic to four islands in the Philippine archipelago (Mindanao, Leyte, Samar, and Luzon; Fig. 1), but now sparsely distributed across mainly montane, and some lowland, dipterocarp forests (Salvador & Ibañez 2006; Sutton *et al*. 2023a). The population has declined drastically over the past 50 years, mainly due to habitat loss through deforestation (Kennedy 1977; Bueser *et al*. 2003; Sutton *et al*. 2023a) and persecution (Salvador & Ibañez 2006; Ibañez *et al*. 2016a).

**Figure 1.**
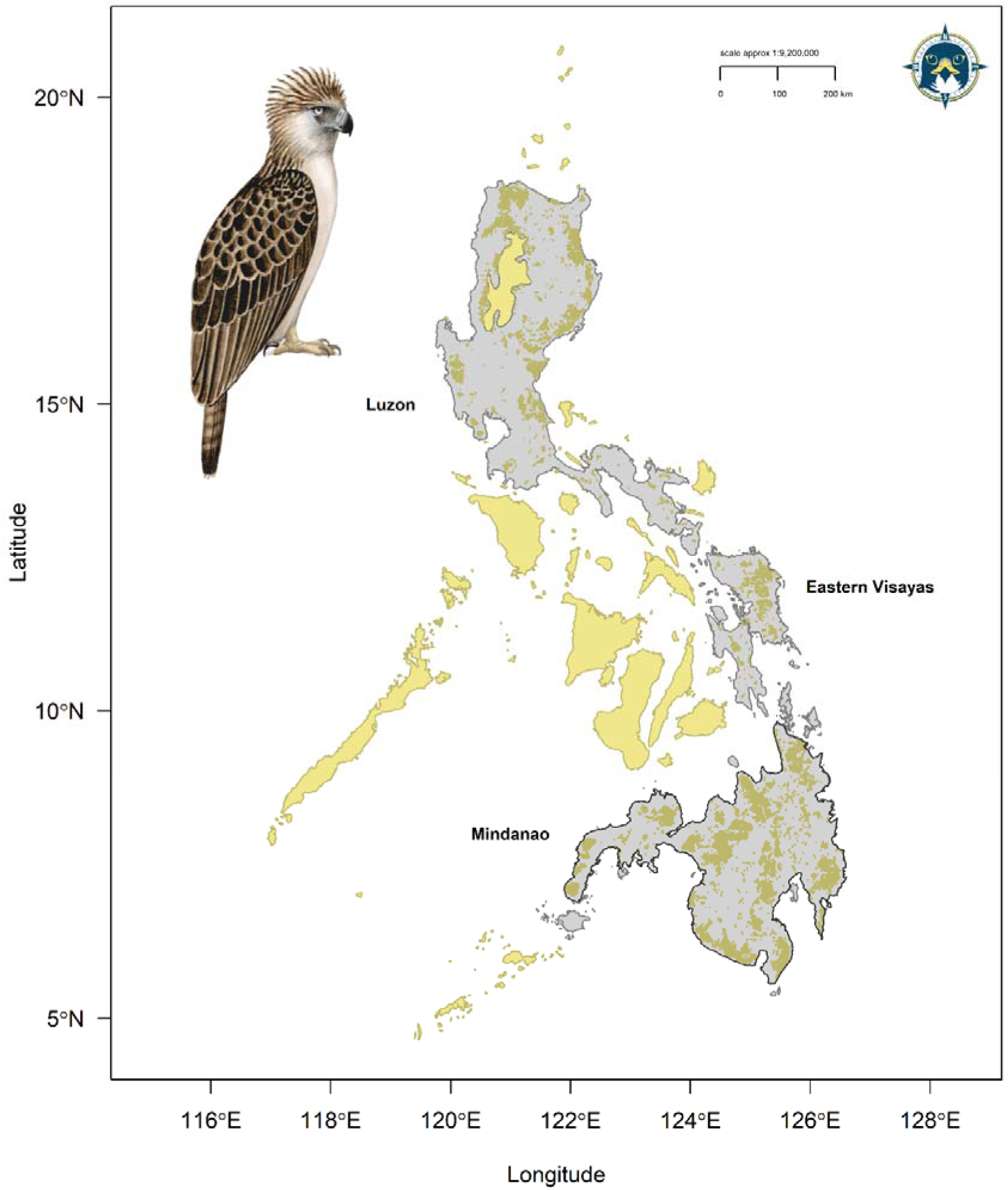
Current distribution map for the Philippine Eagle showing range islands (light grey) and Area of Habitat range model from Sutton *et al*. (2023a; brown polygons). Dark grey polygon defines the extent of Mindanao for our modelling. Yellow polygons define the national boundary of the Philippines outside of the species known range. Illustration by Bryce W. Robinson.

The island of Mindanao is the current stronghold for the species, particularly the east of the island, with the most extensive area of habitat and consequently highest predicted population carrying capacity (Sutton *et al*. 2023a). As a forest-dependent apex predator, the Philippine Eagle is acutely vulnerable to habitat destruction caused by mining because many planned mining concessions coincide spatially with areas of high forest canopy biomass which nesting adult eagles select preferentially (Sutton *et al*. 2024a). Thus, mining activities have the potential to destroy nest sites, reduce prey availability, and limit breeding success (BirdLife International 2018). Therefore, understanding where mining concessions are concentrated, and to what extent Philippine Eagle nest habitats may be impacted by mining operations is critical for setting spatial conservation priorities for the species.

Spatial gap analysis has traditionally been used to evaluate the coverage of protected areas against the distribution of suitable habitats (Scott *et al*. 1993; Sutton *et al*. 2023b). For the Philippine Eagle, this analysis has revealed discrepancies between the eagle’s habitat needs and the protection provided by existing reserves (Sutton *et al*. 2023a). Such gaps in protection often occur in regions rich in mineral resources, where mining pressures are high. Hotspot analysis is a complementary approach to gap analysis which then identifies locations with a high concentration of suitable habitats (Erst *et al*. 2023). A spatial gap analysis integrated with a hotspot analysis assessed with a predictive species distribution model (SDM) offers a promising approach to mitigating the impacts of mining on the Philippine Eagle. For the Philippine Eagle, SDMs can identify suitable habitats by incorporating ecological variables such as forest cover and topography, similar to previous studies quantifying landscape-scale habitat use for large forest raptors (Sutton *et al*. 2022a; Natsukawa *et al*. 2024).

By mapping areas of high habitat suitability, SDMs help pinpoint critical areas that overlap with existing or planned mining concessions. This information is crucial for highlighting high-risk zones where mining activities could significantly threaten eagle populations. Additionally, understanding the current coverage of both protected areas and indigenous land areas is key for Philippine Eagle conservation (Salvador & Ibañez 2006). Many indigenous lands are in montane forest remnants where most Philippine Eagles nest (Sutton *et al*. 2023a). Indigenous communities hold the eagle in high regard and seek to protect habitat for the species; thus these indigenous areas and communities are crucial for eagle conservation (Salvador & Ibañez 2006; Panopio *et al*. 2021). Importantly, many planned mining concessions are within areas of indigenous managed land (known as ‘ancestral domains’), so understanding the extent of habitat across ancestral indigenous areas is key for Philippine Eagle conservation priority setting (Panopio *et al*. 2021).

Nest surveys are a crucial tool for assessing the impact of mining on wildlife, especially for species like the Philippine Eagle that require specific conditions for nesting and reproduction (Abaño *et al*. 2016; Salvador & Ibañez 2006). By identifying nesting sites and their proximity to mining activities based on a predictive SDM, researchers can evaluate the extent of habitat disturbance and prioritize areas for conservation (Ibañez *et al*. 2003). This information can guide the creation of buffer zones around critical habitats and inform land-use planning to minimize overlap between mining operations and key nesting areas. For species like the Philippine Eagle, these surveys are invaluable in shaping conservation strategies that mitigate the impacts of mining while ensuring the protection of the species’ remaining habitats (DENR 2016).

Here, we aim to: (**1**) model the current habitat distribution of the Philippine Eagle on Mindanao in relation to environmental variables and known occurrences; (**2**) analyse the overlap between suitable eagle habitats and existing or planned mining concessions to identify high-risk zones within the EMBC; and (**3**) evaluate the effectiveness of existing protected areas and ancestral domains in conserving the Philippine Eagle within the corridor and propose conservation strategies to mitigate mining impacts. We focused our gap and hotspot modelling on the EMBC, which is a global priority for biodiversity conservation having the largest concentration of Key Biodiversity Areas (KBA) on Mindanao, on one hand, and the greatest number of mining concessions on the other (Ambal *et al*. 2012). By addressing these objectives, we seek to provide an evidence base for conservation planning and policy recommendations to balance resource extraction with the conservation of the Philippine Eagle and its habitat.

## Methods

### Species locations

We compiled Philippine Eagle point localities from the Global Raptor Impact Network (GRIN, McClure *et al*. 2021), a data information system for population monitoring of all raptor species. For the Philippine Eagle, GRIN includes presence-only data consisting of nest locations (*n* = 48) from unstructured surveys (i.e., with no true absence data) conducted on Mindanao by the Philippine Eagle Foundation since 1978 to the present (Miranda *et al*. 2000; Ibañez *et al*. 2016a). We used all nest locations recorded from year 1978 onwards because they matched the temporal timeframe and 1-km resolution of the habitat covariates, whilst retaining sufficient sample size for robust modelling (van Proosdij *et al*. 2016). We used a 1-km spatial resolution because this is the standard resolution to calculate the area of habitat and is also an effective area size to conduct nest surveys.

Additionally, we included 562 GPS tracking fixes from seven nesting adult Philippine Eagles on the island of Mindanao sourced from the Philippine Eagle Foundation (Table S1) and pooled this with the nest locations to better represent habitat use of a rare species with limited occurrences (Fletcher *et al*. 2019; See Supplementary Material). A total of 80,966 raw unfiltered fixes were obtained from four adult females and three adult males from April 2013 to September 2024. We removed all duplicate fixes and applied a 1-km spatial filter to this raw dataset resulting in 562 spatially filtered GPS fixes (Table S1). We combined the known nest locations with the filtered GPS tracking fixes, resulting in a filtered subset of 610 occurrence records for the calibration models. We used spatial filtering because it is the most effective method to account for sampling bias (Kramer-Schadt *et al*. 2013; Boria *et al*. 2014; Fourcade *et al*. 2014) and to ensure we retained the nest locations as priority data points because of their geolocation accuracy and direct relevance to optimal conditions and resources for Philippine Eagle nest occurrence.

### Habitat covariates

We defined the species’ accessible area (Barve *et al*. 2011) as consisting of the mainland area of Mindanao (Fig. 1; dark bordered grey polygon), extracted from the World Wildlife Fund (WWF) terrestrial ecoregions shapefile (Olson *et al*. 2001). We selected the ecoregion area corresponding to either lowland or montane moist tropical forest because the Philippine Eagle is a habitat specialist of tropical dipterocarp forests (Bueser *et al*. 2003; Salvador & Ibañez 2006; Sutton *et al*. 2023a). All raster environmental layers were then cropped to this delimited polygon consisting of the ecoregions within the mainland area of Mindanao. We selected habitat covariates *a prioiri* based both on environmental factors related empirically to resources and conditions influencing Philippine Eagle distribution (Bueser *et al*. 2003; Ibañez *et al*. 2003; Salvador & Ibañez 2006; Sutton *et al*. 2023a).

We predicted occurrence using six continuous covariates at a spatial resolution of 1-km (Fig. S1) derived from multiple satellite remote sensing products. We used three forest structure indices to assess the combined effects of forest structure and human impact on the distribution of the Philippine Eagle: Forest Landscape Integrity Index (FLII; Grantham *et al*. 2020), Forest Structural Condition Index (FSCI; Hansen *et al*. 2019) and Forest Structure Integrity Index (FSII; Hansen *et al*. 2019), all sourced from a globally available repository (https://map.unbiodiversitylab.org/earth). We included three indices of forest structure and integrity because they are important predictors for avian SDMs, albeit rarely used (Leitão & Santos 2019; Burns *et al*. 2020), and knowing that low level human infrastructure is empirically related to highly suitable Philippine Eagle habitat (Sutton *et al*. 2023a). Additionally, these indices provide complementary measures of forest condition, ranging from broad landscape integrity to detailed structural attributes, allowing for a comprehensive evaluation of habitat suitability. We extracted FLII values at 300-m resolution and resampled these pixels to 1-km using bilinear interpolation. We utilized FSCI and FSII data at 30-m resolution and then resampled the raster layers to 1-km using bilinear interpolation to refine our analysis of fine-scale habitat quality.

Forest Landscape Integrity Index (FLII; Grantham *et al*. 2020) was utilized to quantify the degree of anthropogenic disturbance at a landscape scale. The FLII integrates measures of human modification, including land-use change, infrastructure, and direct forest exploitation, to generate a continuous index of forest integrity, with higher values indicating minimally disturbed forests. Forest Structural Condition Index (FSCI; Hansen *et al*. 2019) was employed to evaluate forest structure based on remote sensing-derived tree cover, canopy height, and biomass. This index provides an assessment of forest degradation by measuring deviations from expected structural conditions under minimal human disturbance, with higher values indicating less disturbed forests. Last, the Forest Structure Integrity Index (FSII; Hansen *et al*. 2019) was used to measure structural degradation by comparing observed forest structure against a baseline of intact conditions. FSII captures the extent to which human activity has altered vertical and horizontal forest complexity, providing a critical parameter for species reliant on undisturbed forest canopies, with higher FSII values indicating more complex forest structure.

Additionally, we used a Homogeneity texture measure calculated from a 1-km resolution Enhanced Vegetation Index (EVI) and a 1-km Leaf Area Index (LAI) measuring forest canopy leaf density, both sourced from the Dynamic Habitat Indices portal (Radeloff *et al*. 2019; https://silvis.forest.wisc.edu/data/dhis/). Homogeneity is a biophysical texture measure closely related to vegetation structure, composition, and diversity (i.e., species richness) derived from textural features of Enhanced Vegetation Index (EVI) between adjacent pixels; sourced from MODIS vegetation products, spanning the years 2003-2014 (Radeloff 2019). We calculated raster cell texture values using 32 grey-level co-occurrence matrices in the R package glcm (Zvoleff 2019) using a 3x3 window and a 90° shift to represent the spatial variability and arrangement of vegetation species richness on a continuous scale which varies between zero (minimum homogeneity, high species richness) and one (maximum homogeneity, low species richness).

Leaf Area Index (LAI) is a biophysical measure of the amount of foliage within the plant canopy based on MODIS vegetation products, used here as a composite Dynamic Habitat Index product spanning the years 2003-2014 (Radeloff 2019). LAI values range from 0 (bare ground) to > 10 (dense coniferous forest) and is a key driver of primary productivity (Asner *et al*. 2003). The DHI products summarise three measures of vegetation productivity: annual cumulative, minimum throughout the year, and seasonality as the annual coefficient of variation. Combined, we used both Dynamic Habitat Indices as environmental proxies, with higher LAI values associated with higher prey species richness (Hobi *et al*. 2017) and higher EVI values with higher habitat heterogeneity, both of which are key variables for predicting the distribution of large tropical forest eagles (Sutton *et al*. 2023c; Sutton *et al*. 2024b).

Last, we included Terrain Roughness Index (TRI) as a measure of topographic complexity sourced from the European Space Agency Copernicus 30m Global Digital Elevation Model (ESA 2024). TRI data were extracted at 30m resolution from the Open Topography repository (https://portal.opentopography.org) and then resampled using bilinear interpolation to 1-km to match the existing raster layers. TRI is a measure of variation in topography around a central pixel, with lower values indicating flat terrain and higher values indicating larger differences in elevation of neighbouring pixels (Wilson *et al*. 2007). We used TRI because of the complex mountainous landscape of Mindanao, knowing that Philippine Eagles prefer areas of medium to high topographic complexity for nest sites (Ibañez *et al*. 2003). All selected covariates showed low collinearity with Variance Inflation Factors (VIF) all < 6; Table S2).

### Species Distribution Models

We parametrized the SDMs using a fine 1-km pixel grid, equivalent to fitting an inhomogeneous Poisson process (IPP) with loglinear intensity (Baddeley *et al*. 2010). We did this because the IPP framework is the most effective method to model presence-only data (Warton & Shepherd 2010), common to many raptor monitoring programmes which solely seek to identify occupied areas (Geary *et al*. 2018). We fitted SDMs using penalized logistic regression, via maximum penalized likelihood estimation (Hefley & Hooten 2015) in the R package maxnet (Phillips *et al*. 2017). Penalized logistic regression imposes a regularization penalty on the model coefficients, shrinking towards zero the coefficients of covariates that contribute the least to the model, reducing model complexity (Gastón & García-Viñas 2011). We limited model complexity because this is necessary when the primary goal is to use SDMs for predictive transferability in space (Helmstetter *et al*. 2020).

The maxnet package fits the SDM as a form of infinitely weighted logistic regression (presence weights = 1, background weights = 100), based on the maximum entropy algorithm, MAXENT (Phillips *et al*. 2017). MAXENT is designed for presence-background SDMs and is mathematically equivalent to estimating the parameters for an IPP (Renner & Warton 2013; Renner *et al*. 2015). We used a tuned penalized logistic regression algorithm because this approach outperforms other SDM algorithms (Valavi *et al*. 2021), including ensemble averaged methods (Hao *et al*. 2020). We evaluated calibration accuracy for the Mindanao model using a random sample of 5150 background points at a recommended 1:10 ratio to the number of occurrence points (Helmstetter *et al*. 2020), to sufficiently sample the background calibration environment (Barbet-Massin *et al*. 2012; Guevara *et al*. 2018). Full details on the model parameter settings are outlined in the Supplementary Material.

We used Continuous Boyce index (CBI; Hirzel *et al*. 2006) as a threshold-independent metric of how predictions differ from a random distribution of observed presences (Boyce *et al*. 2002). CBI is consistent with a Spearman correlation (*r_s_*) and ranges from -1 to +1. Positive values indicate predictions consistent with observed presences, values close to zero suggest no difference with a random model, and negative values indicate areas with frequent presences having low environmental suitability. We calculated mean CBI using five-fold cross-validation on 20 % test data with a moving window for threshold-independence and 101 defined bins in the R package enmSdm (Smith 2019). We further tested the optimal predictions against random expectations using partial Receiver Operating Characteristic ratios (pROC), which estimate model performance by giving precedence to omission errors over commission errors (Peterson *et al*. 2008). Partial ROC ratios range from 0 to 2 with 1 indicating a random model. Function parameters were set with a 10% omission error rate, and 1000 bootstrap replicates on 50% test data to determine significant (*α* = 0.05) pROC values >1.0 in the R package ENMGadgets (Barve & Barve, 2013).

### Hotspot Analysis

To further refine our analysis, we projected our range-wide continuous SDM for Mindanao into the area of eastern Mindanao (including the provinces of Agusan del Norte and del Sur, Compostela Valley, Davao Oriental and Surigao del Norte and del Sur), to predict hotspots of Philippine Eagle habitat. The projected continuous prediction for eastern Mindanao was then reclassified to a threshold prediction using three discrete quantile classes representing habitat suitability (Low: 0.000 - 0.382; Medium: 0.383 - 0.652; High: 0.653 - 1.000). We calculated class-level landscape metrics on the discrete quantile classes to estimate total and core area of habitat in each class in the R package SDMTools (VanDerWal *et al*. 2014), based on the FRAGSTATS program (McGarigal *et al*. 2002). Core areas were defined as those cells with edges wholly within each habitat class, with cells containing at least one adjacent edge to another class cell considered as edge habitat (Sutton *et al*. 2022b). We then overlaid the current mining concessions shapefile for Mindanao and identified those mining concessions that had continuous core areas of 9 km^2^ of high habitat cells as priority sites for nest surveys. We used 9 km^2^ because this is the minimum core home range estimate for adult Philippine Eagles on Mindanao (Sutton *et al*. 2024a).

Additionally, we validated our models in conjunction with expert judgement because this approach gives most benefit to conservation priority assessments (Marcer *et al*. 2013; Syfert *et al*. 2014). We assessed the discrete quantile thresholds for biological realism (25% lower quantile, median, 75 % upper quantile), using expert critical feedback to assess the predictive ability of our models, with the 75 % upper quantile discrete class evaluated as a plausible range extent for high habitat suitability. We followed this participatory modelling process methodology to ensure a robust expert validation of our models, concurring with current knowledge of species biology and its application to conservation planning (Ferraz *et al*. 2020).

### Spatial Gap Analysis

Last, we assessed the level of protected area coverage within the Eastern Mindanao Biodiversity Corridor (EMBC) and indigenous ancestral domains using the Philippines Department of Environment and Natural Resources (DENR) terrestrial shapefiles. We overlaid and intersected the EMBC and ancestral domain shapefiles to assess the current level of protected area coverage and identify any key gaps in coverage between the two shapefiles compared to areas of highest core nest habitat. We used the R program (v3.5.1; R Core Team, 2018) for model development and geospatial analysis using the raster (Hijmans 2017), rgdal (Bivand *et al*. 2019), rgeos (Bivand & Rundle 2019) and sp (Bivand *et al*. 2013) packages.

## Results

### Species Distribution Models

From our 38 candidate models, only one had an ΔAIC_c_ ≤ 2 with this best-fit model using a beta coefficient penalty of β = 1 with both linear and quadratic terms as model parameters. Our best-fit model had high calibration accuracy (mean CBI = 0.936), and was robust against random expectations (pROC = 1.602, SD± 0.049, range: 1.427 – 1.751). The optimal model shrinkage penalty (β = 1) was able to retain eleven non-zero beta coefficients (Table 1), meaning most covariate terms were highly informative to model prediction (Figs. S4-S6). Philippine Eagles on Mindanao were most strongly associated with low Homogeneity values, meaning that high habitat heterogeneity derived from Enhanced Vegetation Index was the optimal predictor for identifying high-suitability eagle nest habitat (Table 1; Fig. 2).

**Figure 2.**
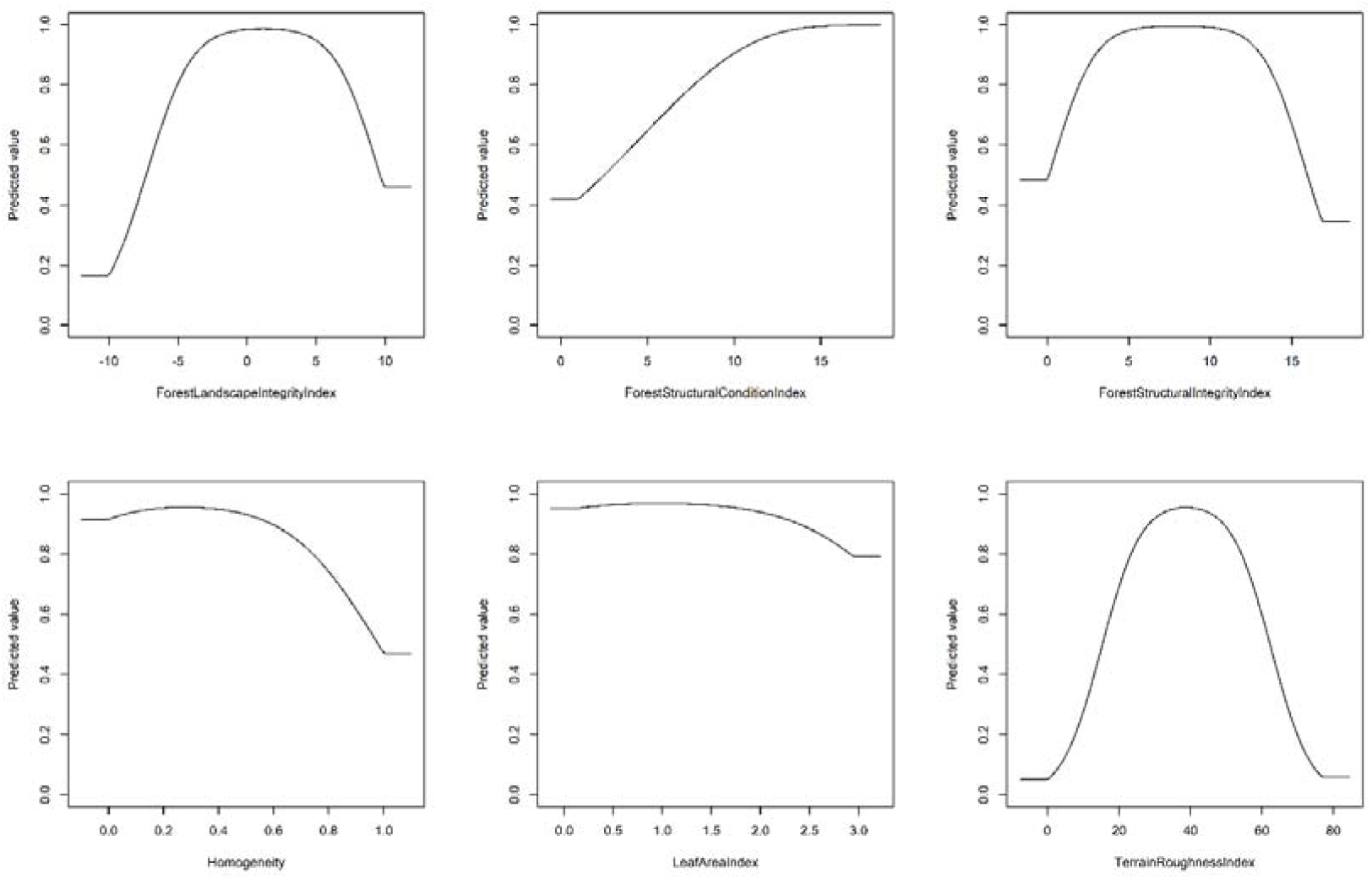
Penalized logistic regression response functions for each habitat covariate from the optimal Species Distribution Model for the Philippine Eagle on Mindanao island. Each curve shows the contribution to model prediction (y-axis) as a function of each continuous habitat covariate (x-axis). Maximum values in each response curve define the highest predicted relative suitability. The response curves reflect the partial dependence on predicted suitability for each covariate and the dependencies produced by interactions between the selected covariate and all other covariates.

**Table 1.** Parameter estimates from penalized linear and quadratic beta coefficients derived from the response functions for each habitat covariate from the range-wide Species Distribution Model for the Philippine Eagle on Mindanao.

| Habitat Covariate | Parameter estimate |  |
| --- | --- | --- |
|  | Linear | Quadratic |
| Forest Landscape Integrity Index | 0.059 | -0.025 |
| Forest Structural Condition Index | 0.163 | 0.000 |
| Forest Structural Integrity Index | 0.523 | -0.033 |
| Homogeneity | 1.687 | -3.045 |
| Leaf Area Index | 0.393 | -0.204 |
| Terrain Roughness Index | 0.212 | -0.003 |

Forest Structural Integrity Index (FSII) had a strong unimodal association, with FSII values ranging from four to 11 most suitable for predicting eagle nest habitat (Table 1). Leaf Area Index (LAI) was the next strongest linear predictor, with LAI values ranging from 1.0 to 2.0, followed by a unimodal relationship with Terrain Roughness Index values peaking at 40, meaning eagle nest habitat was positively associated with medium density leaf canopy cover in areas of low to medium terrain complexity (Fig. 2). Forest Structural Condition Index values had a clear linear association, with values peaking at 15, with Forest Landscape Integrity Index (FLII) values plateauing either side of zero, meaning eagle nest habitat had a positive linear relationship with areas of high forest structure (**ß** = 0.163), and a unimodal relationship with forest landscape integrity (Table 1; Fig. 2).

The largest continuous areas of Philippine Eagle nest habitat across Mindanao were confined to mountainous regions with high forest cover across the eastern and central mountain ranges of Kitanglad, Pantaron, Diwata, and the Bukidnon plateau (Fig. 3). Patchy areas of habitat were identified throughout western Mindanao, largely confined to areas of steep, forested terrain, and extending further south into the Tiruray Highlands and Mount Latian complex. Little habitat was predicted across the now largely deforested lowland plains. The Eastern Mindanao model was able to capture key areas of habitat when projected to eastern Mindanao (Fig. 4). Most high nest habitat suitability for the Philippine Eagle was predicted within the boundaries of the Eastern Mindanao Biodiversity Corridor (EMBC), except for the central cluster (#4) where a small but significant area was predicted to be suitable nest habitat (Fig. 4).

**Figure 3.**
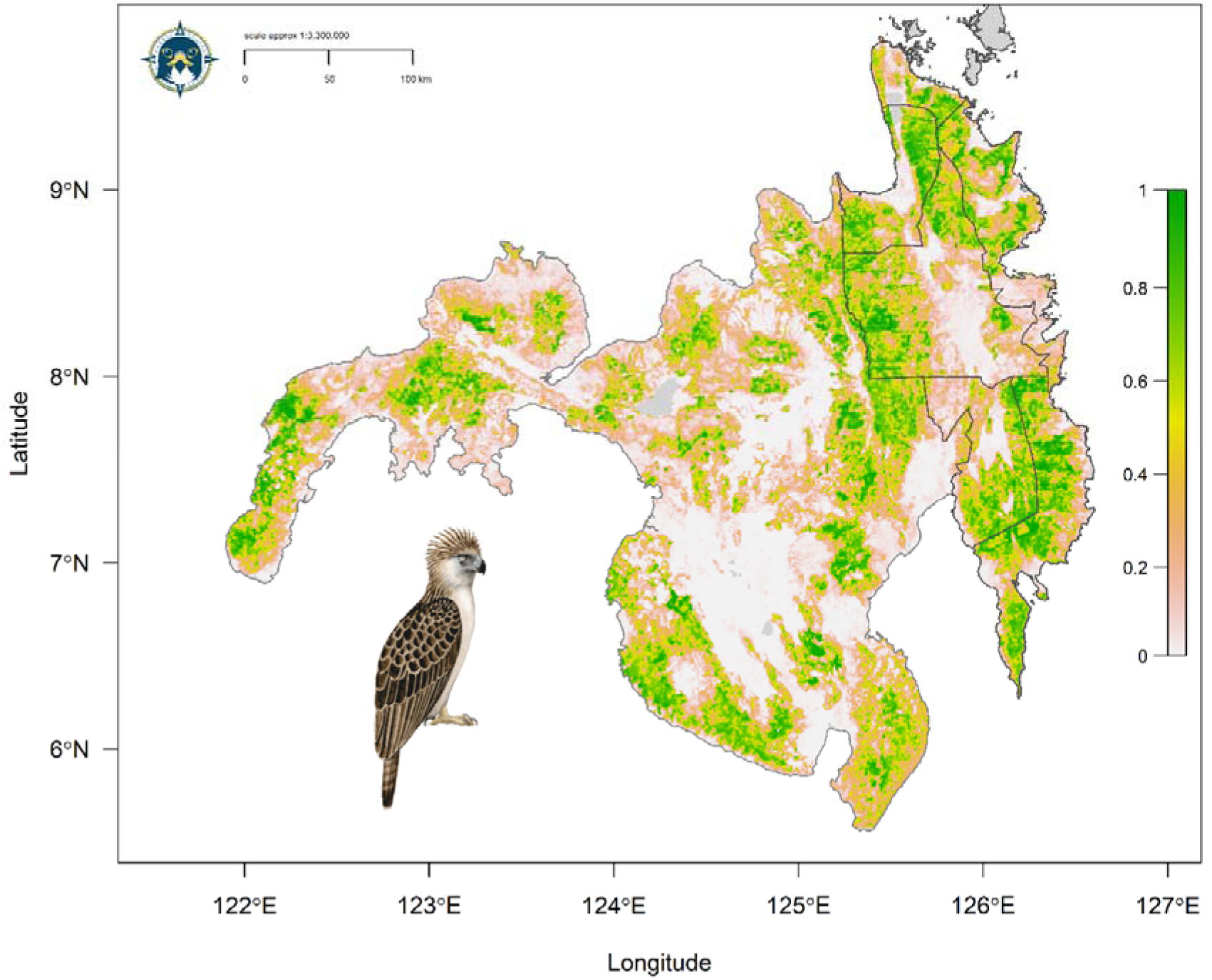
Range-wide Species Distribution Model for the Philippine Eagle on the island of Mindanao using a penalized logistic regression model algorithm. Map denotes habitat suitability prediction with green areas (values closer to 1) having highest habitat suitability, yellow/brown moderate suitability and white low suitability (values closer to zero). Dark grey border defines eastern Mindano region. Illustration by Bryce W. Robinson.

**Figure 4.**
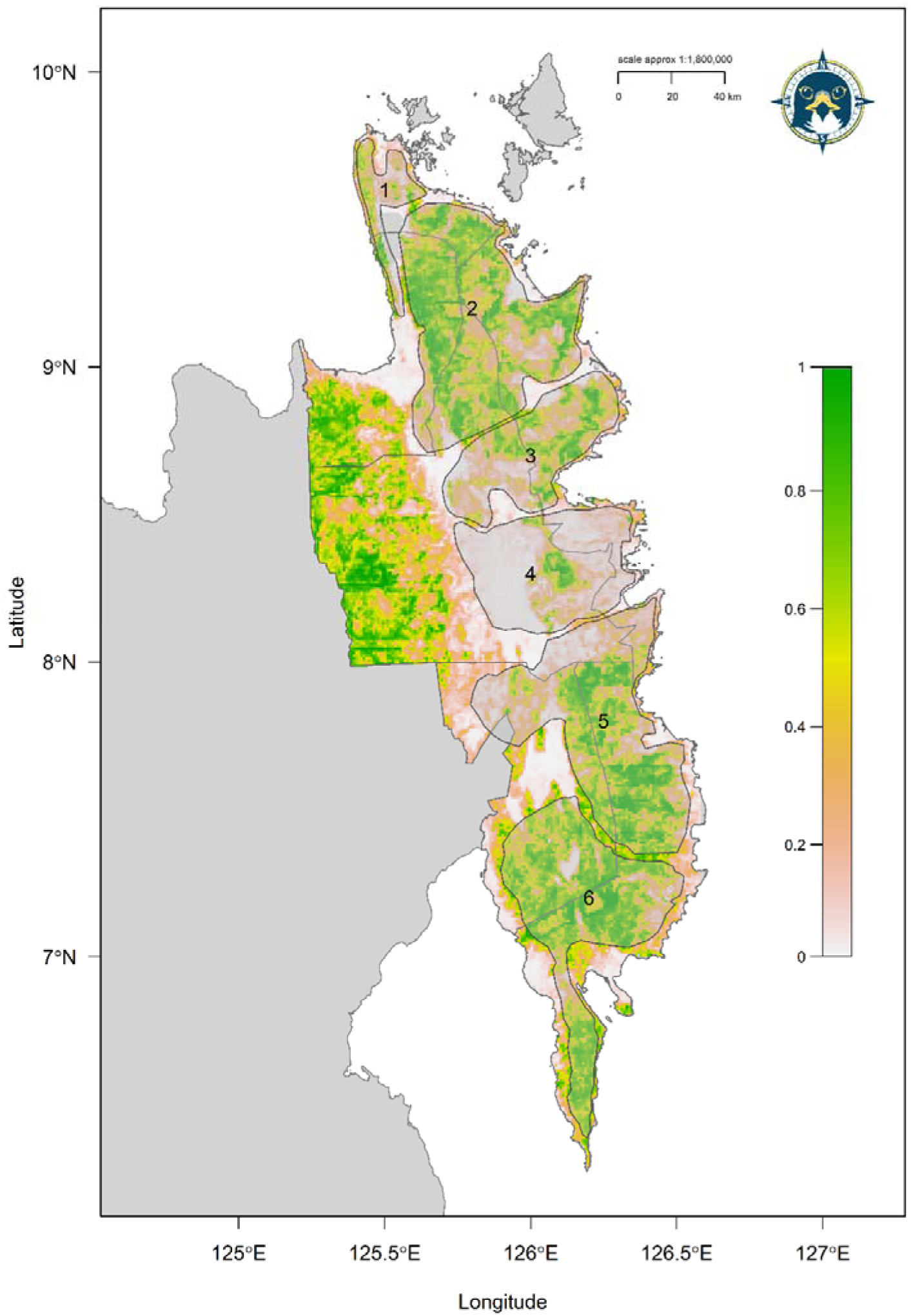
Projected Species Distribution Model for the Philippine Eagle in eastern Mindanao using a penalized logistic regression model algorithm. Map denotes habitat suitability prediction with green areas (values closer to 1) having highest habitat suitability, yellow/brown moderate suitability and white low suitability (values closer to zero). Light grey polygons define Eastern Mindanao Biodiversity Corridor clusters with respective numbers defining each polygon cluster.

### Nest habitat hotspots

The reclassified discrete model for the entire eastern Mindanao region (Fig. 5) calculated nest habitat area in the High quantile class (≥ 0.653) totalling 7 791 km^2^, comprising 29.6 % of the total landscape area of 26 259 km^2^, with 7 999 km^2^ in the Medium quantile class (30.5 %) and 10 469 km^2^ in the Low quantile habitat class (39.9 %). Core areas of High-class nest habitat across eastern Mindanao totalled 2 158 km^2^ of the total core landscape of 8 047 km^2^, comprising 26.8 % of the total High core habitat area, with 475 km^2^ in the Medium class (5.9 %) and 5 414 km^2^ in the Low habitat class (67.3 %). A high proportion of both total (41 %) and core (65 %) High-class nest habitat was predicted to fall within approved mining concessions (Tables 2 & 3; Fig. 5).

**Figure 5.**
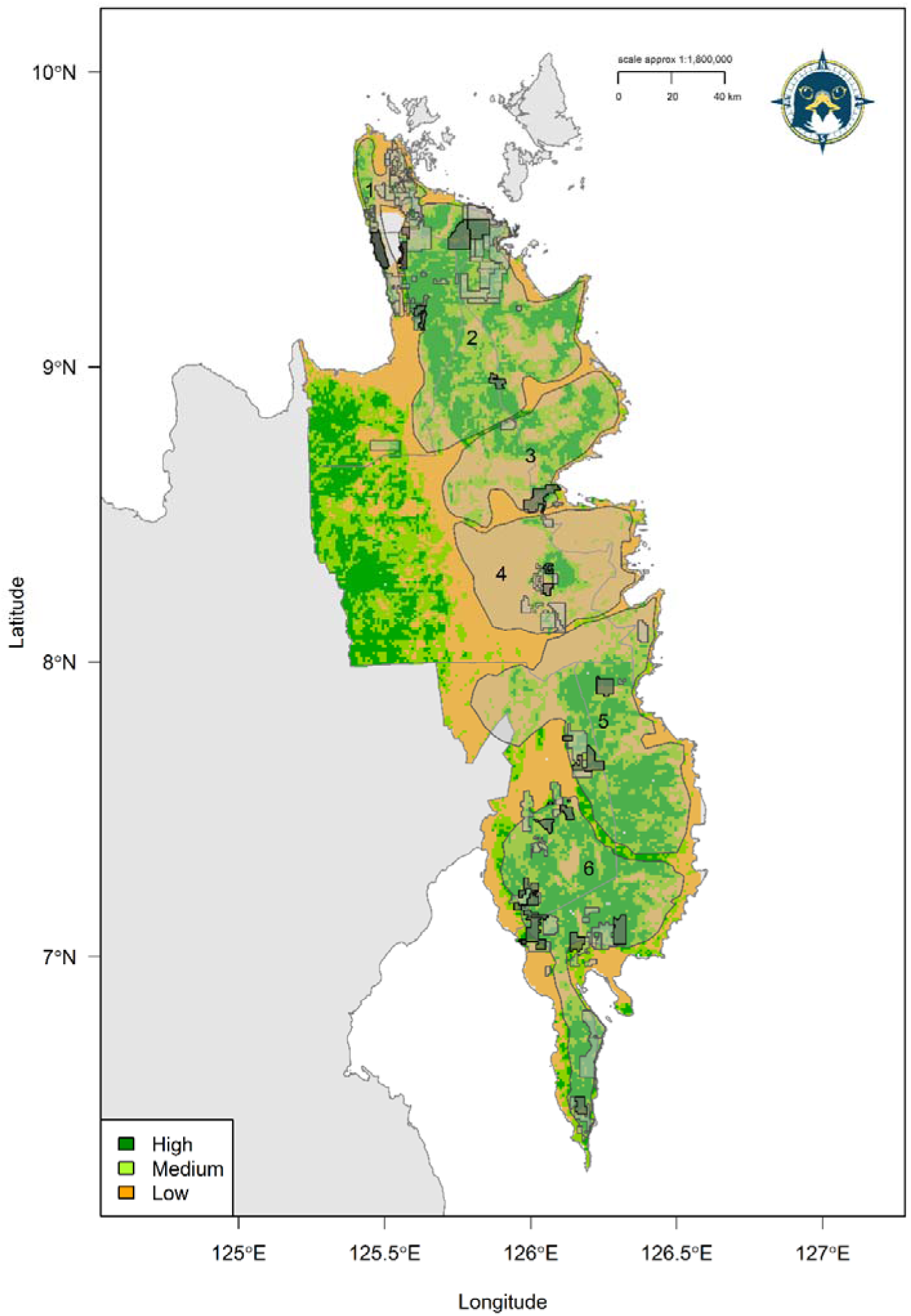
Projected discrete Species Distribution Model for the Philippine Eagle in eastern Mindanao overlaid with priority mining concessions designated for nest surveys (black polygons). Light grey polygons define Eastern Mindanao Biodiversity Corridor clusters with respective numbers defining each polygon cluster.

### Priority conservation areas

Priority mining concessions where resource extraction will intersect with predicted High core nest habitat and should be paused for nest surveys to confirm presence were identified across all six EMBC clusters (Table 4) but with the majority in clusters # 2 and # 6 (Fig. 5). For brevity and visual clarity, the largest mining concessions by area were identified which contain the highest amount of core High-class nest habitat as priority sites for nest surveys and conservation action (Fig. 5). The EMBC protected area clusters covered 35 % of total High-class nest habitat, followed by 31 % and 34 % coverage in the Medium and Low classes respectively (Table 2) but with core High-class habitat coverage greater (40 %; Table 3). Ancestral domain coverage of both total (46 %) and core (65 %) High-class habitat was greater than within the EMBC (Tables 2 & 3; Fig. 6).

**Figure 6.**
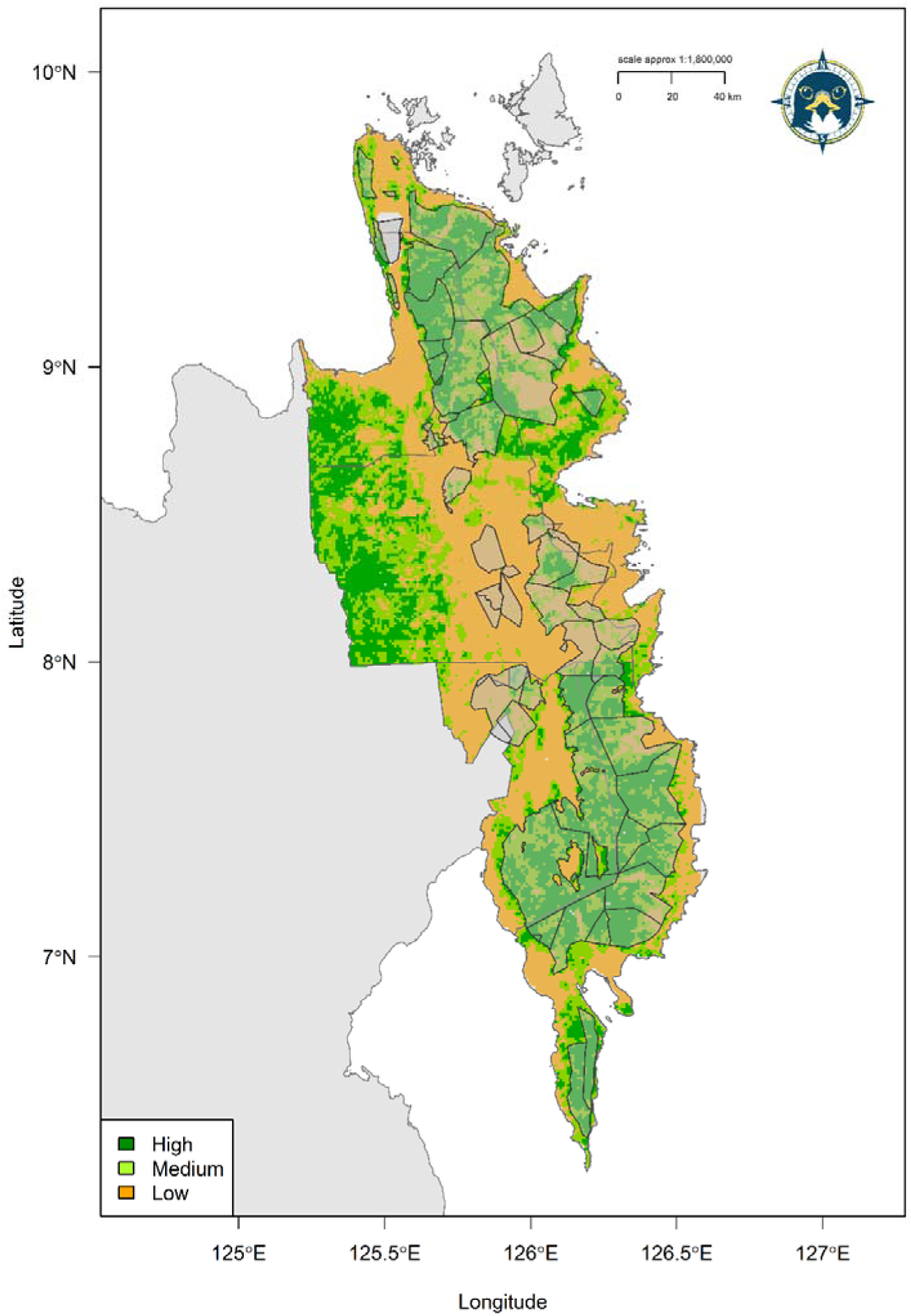
Projected discrete Species Distribution Model for the Philippine Eagle in eastern Mindanao overlaid with indigenous ancestral domains (light grey polygons).

**Table 2.**
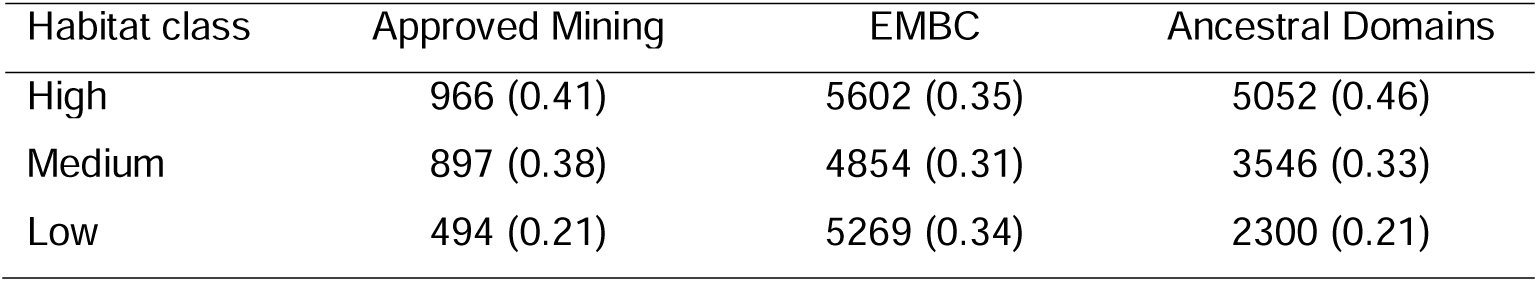
Habitat class landscape metrics for Philippine Eagle nest habitat from the discrete model for eastern Mindanao. Values given are 1-km^2^ grid cells, with their respective proportions in parentheses.

**Table 3.** Habitat class landscape metrics for Philippine Eagle core nest habitat from the discrete model for eastern Mindanao. Values given are 1-km^2^ grid cells, with their respective proportions in parentheses.

| Habitat class | Approved Mining | EMBC | Ancestral Domains |
| --- | --- | --- | --- |
| High | 103 (0.65) | 1702 (0.40) | 1585 (0.65) |
| Medium | 28 (0.18) | 257 (0.06) | 184 (0.07) |
| Low | 26 (0.17) | 2298 (0.54) | 686 (0.28) |

**Table 4.** Priority mining concessions in the Eastern Mindanao Biodiversity Corridor where resource extraction should be paused for nest surveys to be conducted to confirm presence of Philippine Eagles.

| Mining Concession # | Cluster # | Holder | Commodity |
| --- | --- | --- | --- |
| EP-000027-XIII | 1 | Apical Mining Corp. | Gold |
| MPSA-284-2009-XIII SMR | 2 | Kepha Mining Exploration Co. | Nickel, Cobalt, Iron |
| MPSA-007-92-X-SMR | 2 | Suriago Integrated Resources Corp. | Nickel |
| EP-000030-XIII | 2 | SEMCO Exploration and Mining Corp. | Gold |
| MPSA-109-98-XIII | 2 | Consolidated Ores Philippines Inc. | Gold |
| MPSA-344-2010-XIII | 3 | Philex Gold Philippines Inc. | Gold |
| MPSA-299-2009-XIII | 4 | Philsaga Mining Corp. | Gold |
| EP-000003-2010-XI | 5 | Boston Tribal MPurpose Corp. | Gold, Silver |
| EP-000001-2001-XI | 5 | Agusan Petroleum & Mineral Corp. | Gold, Silver, Copper |
| EP-000002-09-XI | 6 | Agusan Metals Corp. | Gold, Silver |
| MPSA-094-97-XI | 6 | Alsons Devt & Investment Corp. | Gold, Silver |
| MPSA-336A-2010-XI | 6 | Napnapan Mineral Resources Inc. | Gold, Silver |
| MPSA-320-2010-XI | 6 | Oro East Mining Co. Inc. | Gold, Silver |
| MPSA-263-2008-XI | 6 | Sinophil Mining and Trading Corp. | Unknown |
| MPSA-197-2004-XI-AMD | 6 | Austral-Asia Link Mining Corp. | Nickel |

## Discussion

By employing a species distribution model in conjunction with spatial gap and hotspot analysis, we identified high-risk zones for a Critically Endangered raptor where mining concessions overlap with predicted nest habitat. Using integrated spatial analysis, we predicted how mining is likely to affect key nesting areas across the species’ range in eastern Mindanao. Our findings indicate that several large mining concessions overlap with predicted high suitability eagle habitat, particularly in regions of high forest cover within ancestral domains. Prioritising those large mining concessions for nest surveys is now paramount to allow surveys to be conducted and mining operations should be paused whilst this is carried out.

### Habitat Use

The predicted habitat suitability map indicates that the largest continuous areas of suitable habitat are concentrated in the Kitanglad, Pantaron, Diwata, and Bukidnon mountain ranges. These montane forests provide nesting and foraging resources, ensuring the ecological resilience necessary for the survival of the Philippine Eagle (Sutton *et al*. 2024a). However, despite the high habitat suitability of these regions, the threat of habitat fragmentation due to mining concessions remains a significant concern. Mining activities in the Philippines accelerate forest loss, leading to severe impacts on biodiversity (Posa *et al*. 2008; Sonter *et al*. 2018). For species like the Philippine Eagle, whose home ranges cover extensive forested landscapes, habitat fragmentation can limit dispersal, reduce gene flow, and increase the risk of local extinctions (Kennedy 1977; Krupa 1989).

### Impact of Mining on Philippine Eagle Habitats

A significant proportion of predicted Philippine Eagle nest habitat (41 % of total and 65 % of core areas) falls within active or proposed mining concessions. This level of habitat encroachment is particularly concerning given the species’ specialized breeding requirements and sensitivity to human disturbance. Mining operations lead to direct habitat destruction through deforestation and introduce secondary threats such as increased human activity, noise pollution, and fragmentation of remaining forest patches (Sonter *et al*. 2018; Laurance *et al*. 2014). These cumulative impacts are especially detrimental to large, territorial raptors like the Philippine Eagle, which require undisturbed forest canopies for nesting and hunting (Miranda *et al*. 2000; Ibañez *et al*. 2016a).

Mining-induced fragmentation also increases the vulnerability of eagle populations to other anthropogenic threats, such as poaching and human-wildlife conflict. For example, fragmented landscapes often facilitate easier access for poachers to eagle territories, leading to an increased risk of illegal hunting and nest disturbance (Gonzales *et al*. 2005; Ibañez *et al*. 2016a). Additionally, the opening of forested areas for mining infrastructure creates new human-wildlife interfaces, increasing the likelihood of retaliatory killings when eagles prey on domestic animals (Paguntalan & Jakosalem 2008). Previous work on the Harpy Eagle (*Harpia harpyja*), another threatened tropical forest raptor, showed that habitat fragmentation and increased human encroachment can lead to a significant decline in reproductive success and increased nest abandonment rates (Miranda *et al*. 2021).

The impact of mining is further exacerbated by its intersection with indigenous ancestral domains, which serve as crucial refugia for the Philippine Eagle.Indigenous communities play an essential role in habitat protection, often acting as the first line of defence against illegal logging and poaching (Panopio *et al*. 2021). However, the encroachment of mining into ancestral lands threatens traditional conservation practices and exacerbates habitat loss. Our results show that 46 % of high-suitability habitat and 65 % of core nest habitat overlap with indigenous ancestral domains, underscoring the need for stronger partnerships between conservation organizations and indigenous communities to safeguard critical eagle habitats (Panopio *et al*. 2021).

### Priority Nest Surveys and Mining Moratoriums

Our findings underscore the need for proactive conservation actions to mitigate the impact of mining on Philippine Eagle populations, and we propose that given the high spatial overlap between eagle habitats and mining concessions, systematic nest surveys should be conducted in priority mining sites before any further mining activities are approved. These surveys will help identify critical nesting areas and inform land-use planning decisions. Nest surveys have proven effective in identifying sensitive eagle nesting sites and can provide early warning signs of population stress, such as reduced breeding success or territory abandonment (Abaño *et al*. 2016; Ibañez *et al*. 2003). Similar nest monitoring programs have been implemented successfully for other threatened raptors, such as the Bonelli’s Eagle (*Aquila fasciata*) in Spain, where regular monitoring of nesting sites has informed conservation actions and reduced nest disturbance (Carrete *et al*. 2002).

In places where mining poses a direct threat to breeding eagle populations, temporary moratoriums on new mining concessions should be enacted while surveys and conservation assessments are conducted. The identification of alternative resource extraction sites that minimize environmental impact should also be explored. Such moratoriums would provide a critical window of opportunity to assess the ecological impact of ongoing mining activities and evaluate the effectiveness of mitigation measures (Sonter *et al*. 2018; WWF 2015). Similar moratoriums have been successful in protecting biodiversity hotspots, such as in the Amazon basin, where restrictions on deforestation for mining have reduced habitat loss and carbon emissions (Finer *et al*. 2013).

Once active nesting sites are identified and delineated, such “critical habitats” should be excluded and legally protected from any form of mining and other extractive resource uses. Philippine Eagles exhibit very high nest site fidelity – generations of breeding pairs use the same nest site repeatedly, which makes keeping nesting sites protected a key factor in the survival and conservation of the species (Ibanez 2009, Ibanez *et al*. 2016b). Additionally, establishing buffer zones around known nesting sites can minimize disturbance from mining activities. These buffer zones should be incorporated into environmental impact assessments and enforced through mining regulations. Buffer zones around key nesting areas can reduce disturbance and enhance reproductive success in sensitive raptor populations (Gonzales *et al*. 2005; Paguntalan & Jakosalem 2008).

We caution however against any interpretation of the absence of Philippine Eagles in high-quality habitat to justify its destruction. Even if current surveys find such areas unoccupied, these habitats are essential for the species’ long-term survival and potential recovery (Sutton *et al*. 2023a). Limiting conservation efforts only to areas currently occupied risks confining the species to its present, often fragmented, distribution, preventing future recolonization or range expansion. High-quality yet unoccupied habitats may serve as crucial dispersal corridors, future breeding grounds, or climate refugia. Destroying habitat simply because it is temporarily unoccupied is short-sighted and incompatible with the goals of long-term species recovery.

### Strengthening Indigenous Land Management

Indigenous communities play a vital role in protecting Philippine Eagle habitats through traditional ecological knowledge and community outreach (Salvador & Ibañez 2006; Ibañez *et al*. 2016a). Conservation initiatives should build on that traditional ecological knowledge and support indigenous-led habitat monitoring and enforcement efforts. Co-management agreements between indigenous groups and government agencies can enhance habitat protection within ancestral lands (Panopio *et al*. 2021). Empowering indigenous communities to manage their ancestral domains not only safeguards eagle habitats but also promotes culturally sustainable conservation practices. Similar initiatives have been successful in protecting Amazonian forests, where indigenous lands have proven more effective in preventing deforestation than strictly protected areas (Nepstad *et al*. 2006).

### Protected Areas and Indigenous Land

Our spatial gap analysis revealed that the Eastern Mindanao Biodiversity Corridor (EMBC) contains 35 % of total high-suitability habitat, particularly in regions where mining activities are most concentrated. Our findings suggest that the implementation of protected areas should be prioritized in regions within indigenous ancestral domains where traditional conservation efforts can be built upon through legal recognition and support (Rodrigues Cazalis 2020; Geldmann *et al*. 2013). This can be achieved through the designation of new conservation zones and cooperation with indigenous-managed conservation areas within ancestral domains. Enforcing protected area policy within the EMBC and integrating them with indigenous ancestral domains through Indigenous and Community Conserved Area (ICCA) management or Other Effective Conservation Measure (OECM) modalities will ensure that critical eagle nesting habitats remain intact (Salvador & Ibañez 2006; Sutton *et al*. 2023a).

Our predictive SDM provides a valuable tool for guiding future conservation planning for the Philippine Eagle on Mindanao. We advise that land-use policies should incorporate species distribution models to identify areas where mining activities should be restricted or modified to reduce their ecological impact and recognise that these model-led policy actions are already being implemented following previous SDM research (Sutton *et al*. 2023a). By implementing targeted conservation actions, including habitat protection, nest monitoring, and community-based conservation initiatives, we can mitigate the adverse effects of mining and ensure the long-term viability of the Philippine Eagle. Future research should focus on refining habitat models through long-term monitoring of eagle populations and assessing the effectiveness of conservation interventions. Additionally, collaborations between conservation organizations, government agencies, and indigenous communities will be critical for achieving sustainable conservation outcomes for the Philippine Eagle and its habitat.

## Acknowledgements

We thank all staff and volunteers from the Philippine Eagle Foundation who conducted fieldwork, including local forest guards, nest wardens and indigenous co-researchers, who contributed data to the GRIN information system. LJS thanks The Peregrine Fund’s Tom Cade Endowment and the Philippine Eagle Foundation for funding and CJWM thanks the M.J. Murdoch Charitable Trust for funding. The PEF would like to thank all of its local government units and Indigenous people’s partners, as well as the Department of Environment and Natural Resources through the Biodiversity Management Bureau and its regional and local offices on Mindanao (DENR Regions 9, 10, 11, 12, and 13). This study is also funded by the DENR-UNDP/GEF Biodiversity Corridor Project, with support from the Global Environment Facility (GEF) and implemented by the Department of Environment and Natural Resources (DENR) and the United Nations Development Programme (UNDP).

## Data Accessibility Statement

The raster and shapefile data that support the findings of this study are openly available on the data repository *figshare* 10.6084/m9.figshare.28911785 Due to confidentiality of nest locations for this critically endangered species we are unable to publicly share our occurrence dataset.

## Author Contribution Statement

LJS conducted the analysis and led the writing with support from all other co-authors. Philippine Eagle Foundation staff provided all occurrence data.

## Supplementary Material

### Methods

#### Species locations

All adult Philippine Eagles were trapped using either a modified Bal-Chatri (Miranda & Ibañez 2006) or a large bownet baited with domestic rabbit (*Oryctolagus cuniculus*). Two eagles were instrumented with solar-powered Global Positioning System-Global System for Mobile Communications (GPS-GSM) transmitters (weight = 70 g; Microwave Telemetry, Inc) while five eagles had battery-powered LC4™ Argos-GPS platform transmitter terminal (PTT) fitted (weight = 105g; Microwave Telemetry, Inc), harnessed with Teflon-coated nylon ribbon backpacks. All tags weighed < 3 % of the body weight for all adults tagged. Tags were programmed to transmit on a 2-hr sampling interval for adults 001F, 002F, 004M, 006F, with adult 003F at 24 hrs and adults 005M and 007M at 2 mins. All birds were marked with aluminium leg bands – the four females with blue bands on their left leg, and the two males with green bands on their right leg. All GPS transmitter harnessing was conducted with a Gratuitous Permit to trap and tag the birds in the presence of a veterinarian as required by the national government of the Philippines.

#### Species Distribution Models

In its original implementation MAXENT imposed a ‘lasso’ (least absolute shrinkage and selection operator) regularization penalty, where only the most significant covariates are retained, with uninformative covariates set at zero. Instead, the maxnet package uses an elastic net penalty (via the glmnet package, Friedman *et al*. 2010) to perform automatic covariate selection (lasso) and continuous shrinkage (ridge regression) simultaneously (Zou & Hastie 2005; Phillips *et al*. 2017), evaluating the contribution of all covariates and shrinking low-contribution coefficients towards zero. Elastic net regularization improves predictive accuracy compared to the lasso, in both simulated and real data examples (Zou & Hastie 2005) and may be viewed as a generalization of the lasso. Overall, penalizing model coefficients reduces model variance, resulting in a regression model that generalizes better (Valavi *et al*. 2021). We parametrized the penalized logistic regression model using infinite weighting (presence weights = 1, background = 100), within the inhomogeneous Poisson process framework because this is the most effective method to model presence-background data as used here (Warton & Shepherd 2010; Hefley & Hooten 2015).

Within the maxnet package the complementary log-log (cloglog) link function was selected as a continuous index of habitat suitability, with 0 = low suitability and 1 = high suitability. Phillips *et al*. (2017) demonstrated the cloglog link is equivalent to an inhomogeneous Poisson process and can be interpreted as a measure of relative occurrence probability proportional to a species potential abundance. Optimal-model selection was based on Akaike’s Information Criterion (Akaike 1974) corrected for small sample sizes (AIC_c_; Hurvich & Tsai 1989), to determine the most parsimonious model from two maxnet parameters: regularization beta multiplier (β; level of coefficient penalty) and feature classes (response functions, Warren & Seifert 2011; Phillips *et al*. 2017). Thirty-eight candidate models of varying complexity were built by conducting a grid search with a range of regularization multipliers from 1 to 10 in 0.5 increments, and two feature classes (response functions: Linear, Quadratic) in all possible combinations using the ‘block’ method of spatial cross-validation (*k =* 5) in the ENMeval package in R (Muscarella *et al*. 2014).

Spatial block cross-validation divides the geographical structure of the data according to latitudinal and longitudinal lines, dividing all occurrences into four spatially independent bins of equal numbers. By binning the geographical structure of test data into blocks, the models are projected onto an evaluation region not included in the calibration process. All occurrence and background test points are assigned to their respective bins dependent on location, thus further reducing spatial auto-correlation between testing and training localities (Muscarella *et al*. 2014). We considered all models with a ΔAIC_c_ < 2 as having strong support (Burnham & Anderson 2004) and selected the model with the lowest ΔAIC_c_ that used both feature classes (Linear, Quadratic) with the highest regularization penalty.

## Supplementary Tables

**Table S1.** GPS metadata for the seven tagged adult Philippine Eagles from the island of Mindanao. Fixes are subsampled from the raw data locations using a 1-km sampling rate interval.

| ID | From | To | Fixes |
| --- | --- | --- | --- |
| 001F | 16/02/2014 | 10/05/2015 | 53 |
| 002F | 22/12/2014 | 20/01/2016 | 48 |
| 003F | 11/04/2013 | 19/02/2014 | 25 |
| 004M | 19/04/2014 | 05/08/2014 | 15 |
| 005M | 08/12/2019 | 03/09/2021 | 95 |
| 006F | 15/10/2019 | 05/06/2021 | 89 |
| 007M | 16/11/2021 | 09/09/2024 | 237 |
| Total |  |  | 562 |

**Table S2.** Multi-collinearity test using stepwise elimination Variance Inflation Factor (VIF) analysis. Covariates with VIF < 6 have low correlation with other covariates, and thus are suitable for inclusion in calibration models when further evaluated for ecological relevance.

| Habitat Covariate | VIF |
| --- | --- |
| Forest Structural Condition Index | 5.32 |
| Forest Landscape Integrity Index | 3.95 |
| Terrain Roughness Index | 1.74 |
| Forest Structural Integrity Index | 1.59 |
| Leaf Area Index | 1.47 |
| Homogeneity | 1.01 |

## Supplementary Figures

**Figure S1.**
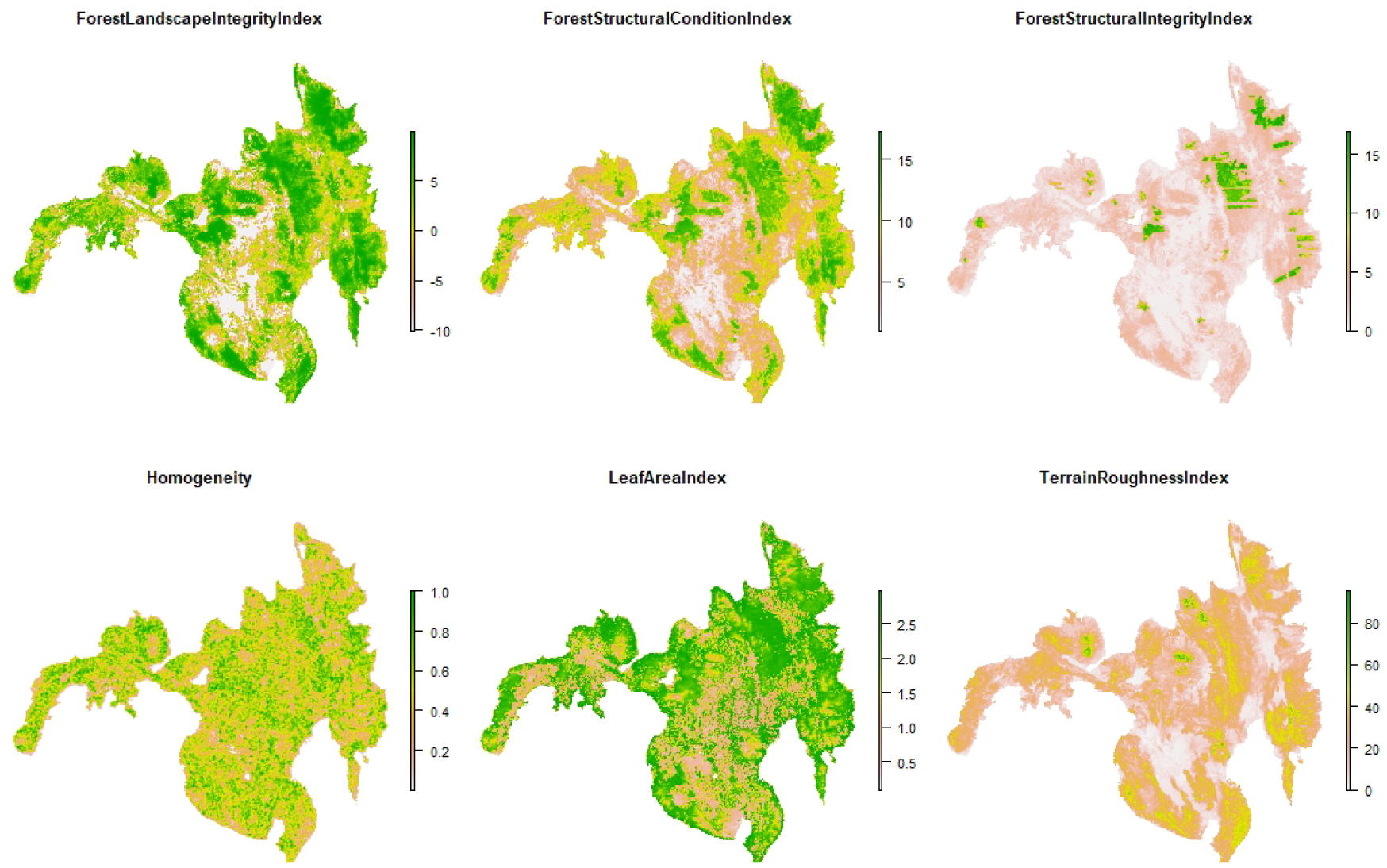
Habitat covariates used in Species Distribution Models for the Philippine Eagle on the island of Mindanao.

**Figure S2.**
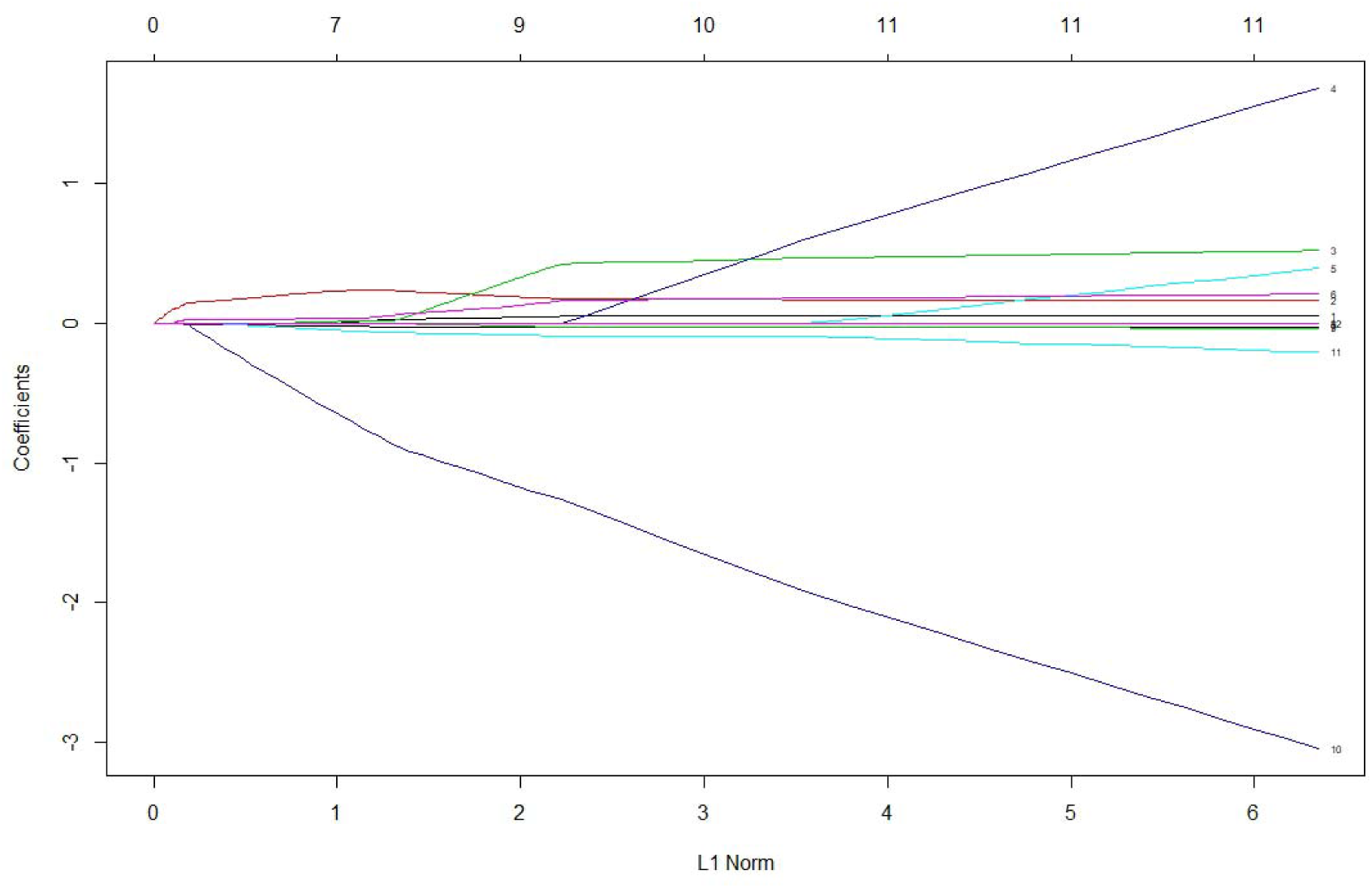
Beta coefficient paths for the optimal penalized logistic regression model where each curve corresponds to a covariate term (linear and quadratic). The paths of each coefficient term are plotted against the L1-norm (lasso or elastic net) of the whole coefficient vector as lambda (the amount defining the level of coefficient shrinkage) varies. The upper axis indicates the number of non-zero coefficients at the current lambda which is the effective degrees of freedom for the lasso or elastic net.

**Figure S3.**
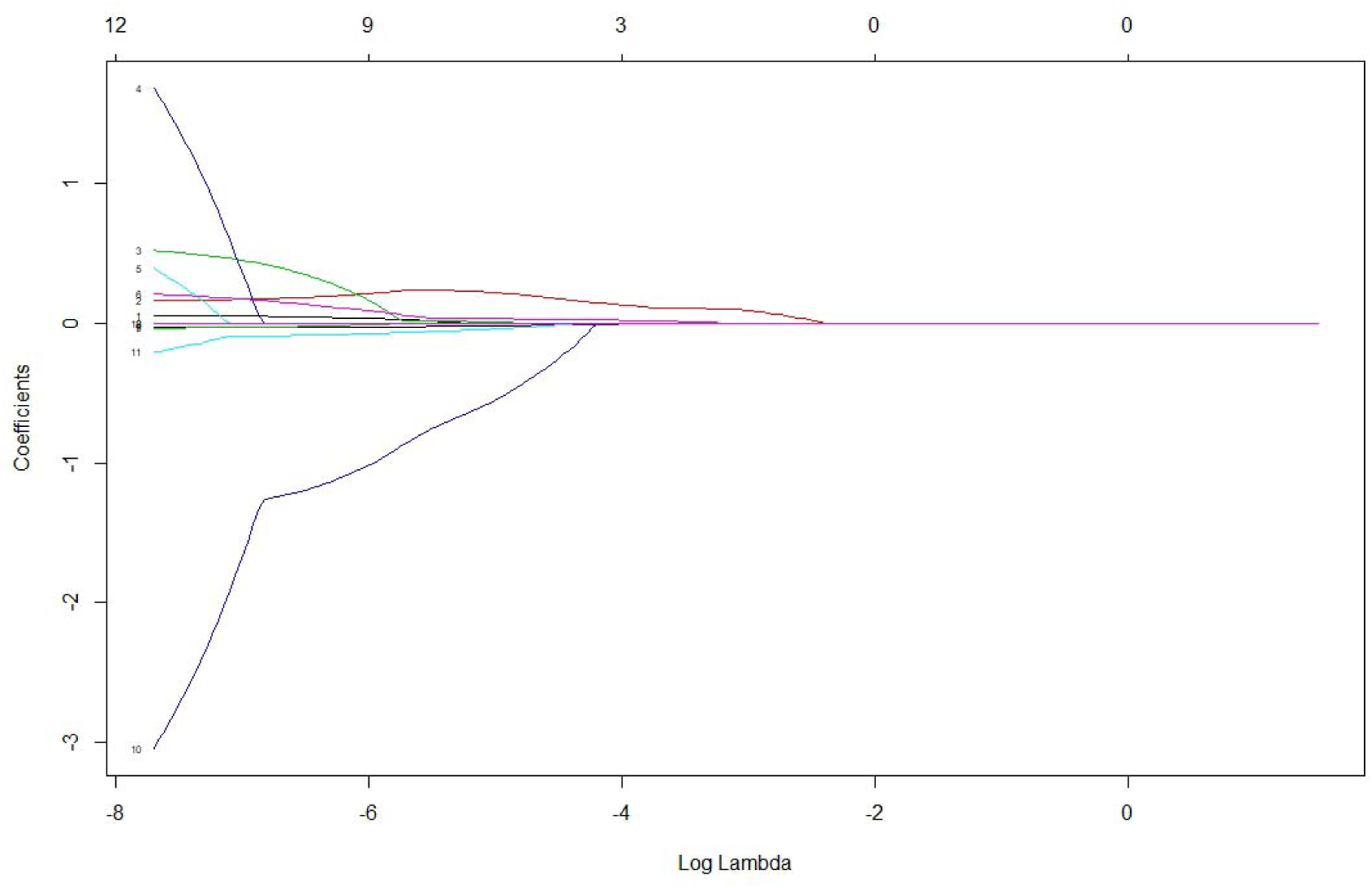
Beta coefficient paths for the optimal penalized logistic regression model where each curve corresponds to a covariate term (linear and quadratic). The paths of each coefficient term are plotted against the log-lambda of the whole coefficient vector as lambda (the amount defining the level of coefficient shrinkage) varies. Log-lambda on the y-axis indicates the log of the optimal value of lambda which minimizes the prediction error. This lambda value will give the most accurate model. The upper axis indicates the number of decreasing non-zero coefficients at the current lambda which is the effective degrees of freedom for the elastic net.

**Figure S4.**
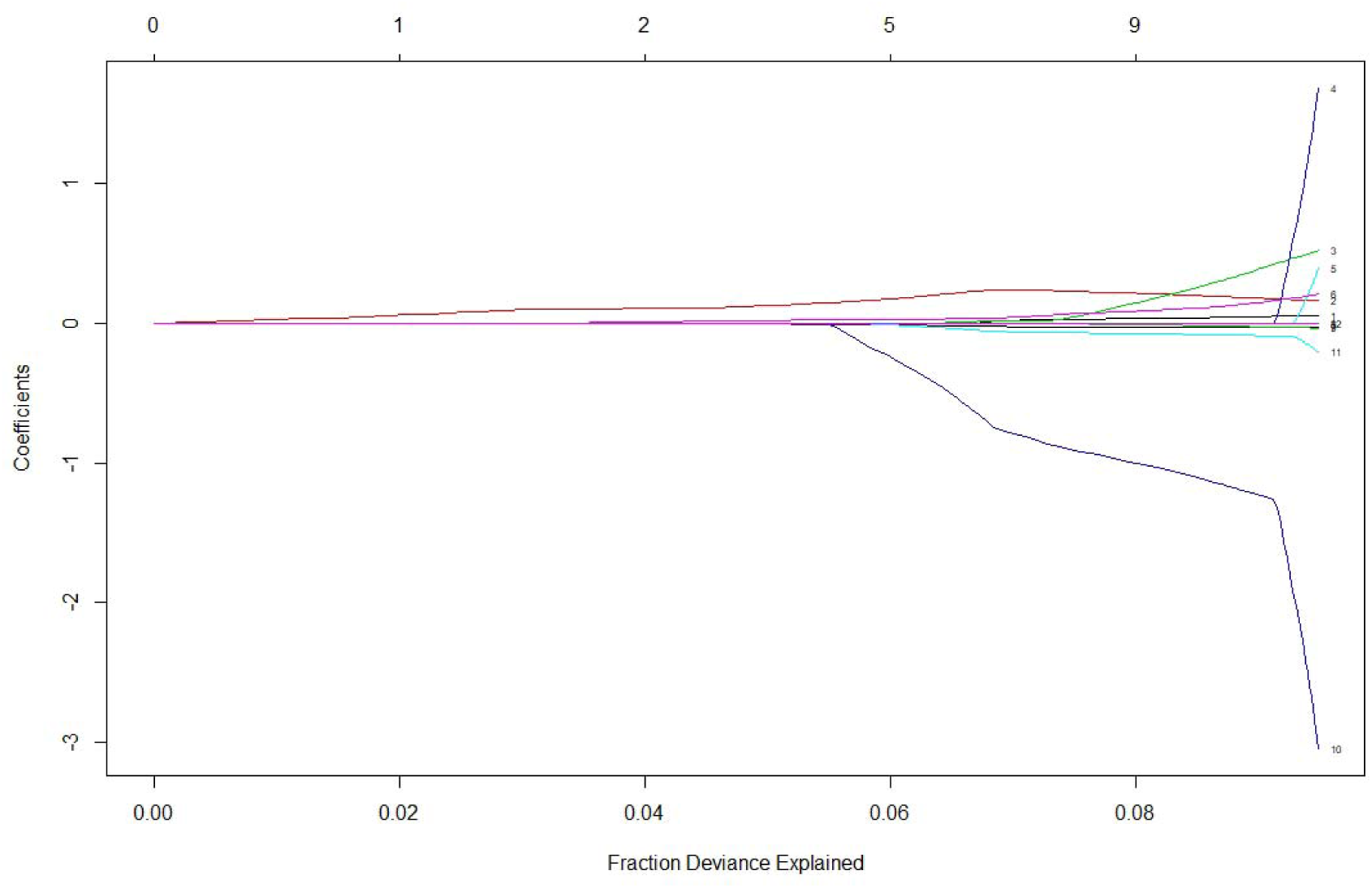
Beta coefficient paths for the optimal penalized logistic regression model where each curve corresponds to a covariate term (linear and quadratic). The paths of each coefficient term are plotted against the fraction deviance explained on the training data. The upper axis indicates the number of non-zero coefficients at the current lambda which is the effective degrees of freedom for the elastic net.

